# An unorthodox mechanism underlying voltage sensitivity of TRPV1 ion channel

**DOI:** 10.1101/2020.01.04.894980

**Authors:** Fan Yang, Lizhen Xu, Bo Hyun Lee, Xian Xiao, Vladimir Yarov-Yarovoy, Jie Zheng

## Abstract

While the capsaicin receptor TRPV1 channel is a polymodal nociceptor for heat, capsaicin, and proton, the channel’s responses to each of these stimuli are profoundly regulated by membrane potential, damping or even prohibiting its response at negative voltages and amplifying its response at positive voltages. Though voltage sensitivity plays an important role is shaping pain responses, how voltage regulates TRPV1 activation remains unknown. Here we showed that the voltage sensitivity of TRPV1 does not originate from the S4 segment like classic voltage-gated ion channels; instead, outer pore acidic residues directly partake in voltage-sensitive activation, with their negative charges collectively constituting the observed gating charges. Voltage-sensitive outer pore movement is titratable by extracellular pH and is allosterically coupled to channel activation, likely by influencing the upper gate in the ion selectivity filter. Elucidating this unorthodox voltage-gating process provides a mechanistic foundation for understanding polymodal gating and opens the door to novel approaches regulating channel activity for pain managements.

---

When the lipid bilayer emerged to enclose living cells more than three billion years ago^1^, this diffusion barrier between cytoplasmic milieu and external environment allowed the establishment of transmembrane ion concentration gradients, which in turn yielded a transmembrane electric potential. Such a membrane potential (V_m_) has been widely utilized in cellular signaling: voltage-gated ion channels alter V_m_ to elicit electric signals for rapid communications^2,3^; voltage-sensitive enzymes, e.g., Ci-VSP^4^, couple changes in V_m_ to the regulation of enzymatic activities and intracellular signaling. For these “classic” voltage-sensing proteins, high sensitivity to voltage (5-fold/mV) has been attributed to a conserved, densely charged “voltage-sensor” domain in the transmembrane region^5^. Other membrane proteins—including G protein-coupled receptors^6^, ion channels and transporters^7^—have also evolved to take cues from V_m_ to perform their biological functions. Understanding the origin and operation of voltage sensitivity in these membrane proteins is of fundamental importance.

Transient receptor potential vanilloid 1 (TRPV1) channel, a polymodal nociceptor in higher species^8^, is a good representative of the non-classic voltage-sensitive membrane proteins. TRPV1 is activated by noxious heat above 40°C^9^, however, its response to heat at depolarized V_m_ is markedly larger than that at the resting V_m_ even when the driving force for its non-selective cation current is equal in magnitude (Fig. 1a). Similarly, TRPV1’s responses to capsaicin and proton—which elicits spiciness sensation^9^ and the sustained phase of pain sensation^10^, respectively—are also substantially asymmetrical (Fig. 1b). Tuning of TRPV1’s nociceptive sensitivity by V_m_ is a highly dynamic process occurring at human body temperature, with the range of voltage dependence (reflected by the conductance-voltage, or G-V, relationship) shifting over a more than 200 mV range in the presence of an activation stimulus^11^ (Fig. 1c). Dynamic voltage sensitivity makes TRPV1 more sensitive to noxious stimuli when its host sensory neuron has been previously excited (primed) or damaged, a general feature for nociception known as hyperalgesia^12^. On the flip side, membrane hyperpolarization is expected to inhibit pain responses, offering opportunities for novel clinical intervention. Despite its physiological significance, how TRPV1 senses V_m_ remains largely elusive.

**Figure 1.**
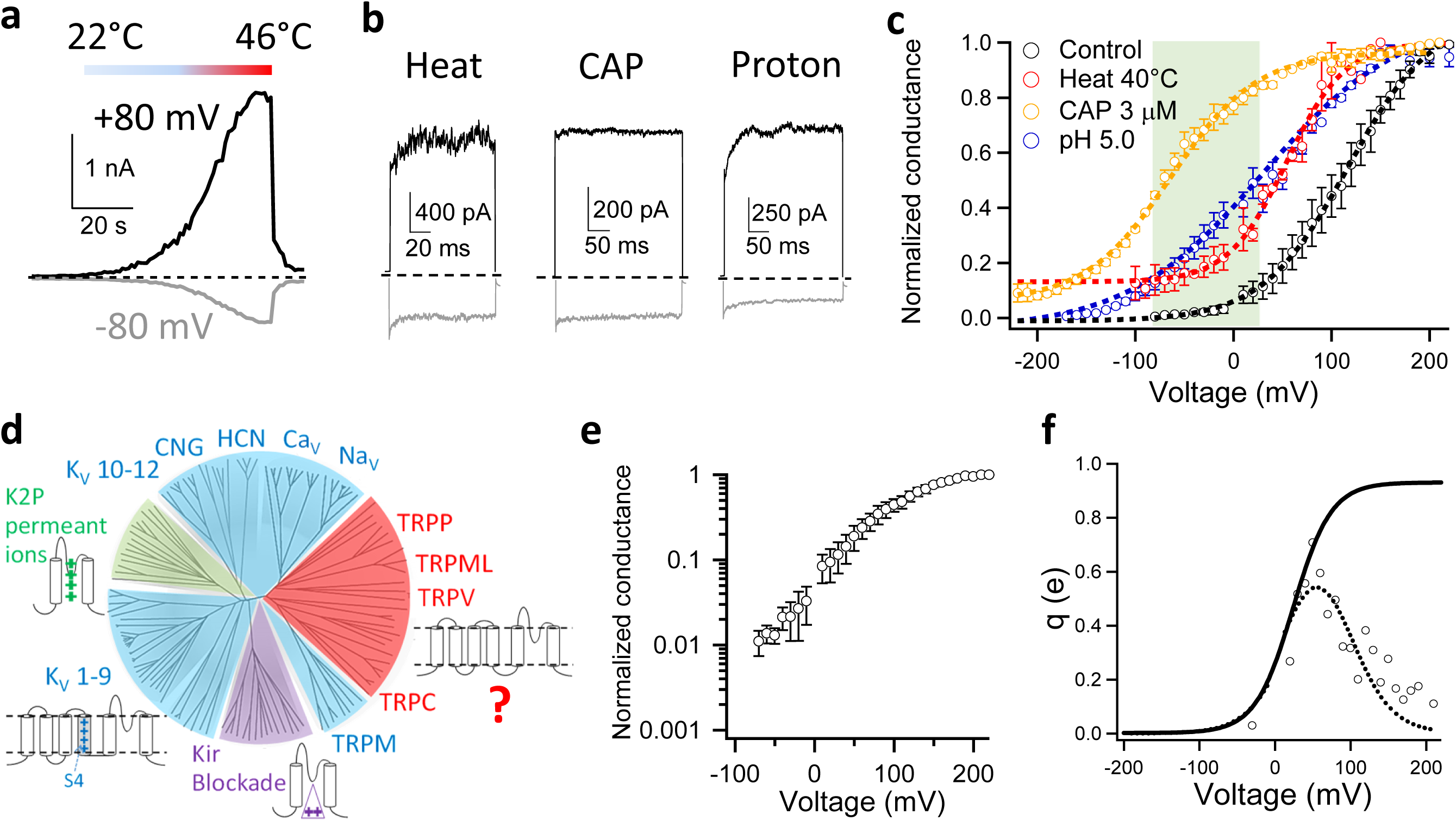
Voltage activation of TRPV1. (**a** and **b**) Transmembrane voltage strongly modulates TRPV1 activities as shown in representative inside-out patch recordings of TRPV1 activated by a heat ramp (**a**), high temperature at 42°C, 1 µM capsaicin (CAP) or pH 5.0 (**b**). Current traces recorded at depolarizing (+80 mV) and hyperpolarizing (−80 mV) voltages are in black and gray, respectively. (**c**) Conductance-voltage (G-V) curve of TRPV1 (black: V_1/2_ = 114.9 ± 1.2 mV, apparent gating charge (q_app_) = 0.72 ± 0.06 e_0_, n = 4) was shifted to more hyperpolarized voltages in the presence of an activation stimulus (red, heat at 40°C: V_1/2_ = 54.8 ± 9.8 mV, q_app_ = 0.94 ± 0.20 e_0_, n = 3; Blue, extracellular pH 5.0: V_1/2_ = 33.4 ± 7.6 mV, q_app_ = 0.46 ± 0.03 e_0_, n = 4; Orange, CAP at 3 µM: V_1/2_ = −72.3 ± 2.3 mV, q_app_ = 0.79 ± 0.19 e_0_, n = 4.). G-V curves are fitted to a single-Boltzmann function (dash curves). Note that with CAP and heat, the G-V curves approach a steady-state level around 0.1 at deeply hyperpolarized voltages. Shaded in green is the physiologically relevant membrane potential range, in which voltage exhibits a strong influence on channel activity. Conductances were normalized to the maximum conductance measured at the highest depolarization voltage. (**d**) A rootless phylogenetic tree representing channels within the VGL ion channel superfamily adapted from a previous report^13^. Sections in blue, green and purple cover channels with a known voltage-sensing mechanism. (**e**) Semi-logarithmic plot of the voltage dependence of normalized mean conductance measured from steady-state current in whole-cell recordings. (**f**) The voltage dependence of the derivative of mean log open probability. Dashed curve is a Boltzmann smoothing function determined from fits to mean log open probability. Solid curve is a fit of the smoothing function with a Boltzmann function at voltages where the open probability is low. All statistics are given as mean ± s.e.m.

TRPV1 belongs to the voltage-gated-like (VGL) ion channel superfamily (Fig. 1d)^13^, for which three voltage-sensing mechanisms have been established. For voltage-gated potassium (Kv), sodium (Nav) and calcium (Cav) channels (blue sections in Fig. 1d), multiple regularly spaced basic residues on the transmembrane S4 helix enable it to move in response to V_m_ changes^14^. For two-pore domain potassium channels (K2P) without a voltage-sensor domain (green section in Fig. 1d), permeant cations have been proposed to bestow voltage sensitivity^15^. For inward-rectifier potassium channels (Kir) (purple section in Fig. 1d), voltage-dependent pore block by endogenous polyamines and Mg^2+^ ions introduces voltage sensitivity^16^. These three mechanisms are applicable to about three-fourth of the VGL superfamily; voltage-sensing is less understood for a significant portion of the family (red section) that is composed of cellular sensor TRP channels including TRPV1. Our study began by determining whether TRPV1 employs one of these three known mechanisms or a new mechanism to detect V_m_.

## Characterizing TRPV1’s voltage sensitivity

A fundamental parameter for voltage sensitivity is the total charge movement, q, which determines the steepness of voltage response^5^. TRPV1 is known to exhibit shallow voltage dependence^11^. By fitting a Boltzmann function to the G-V curve of mouse TRPV1 recorded in HEK293 cells (Fig. 1c), the apparent q was estimated to be 0.72 ± 0.05 e_0_ (n = 4). Applying this approach to Kv channels yielded an apparent q of e_0_^17^. However, it is well known that fitting the shape of G-V curves could yield an under-estimate of q, for example, when there are multiple voltage-dependent closed states that the channel may traverse before reaching the open state^17-19^. Indeed, the total gating charge of Kv, K2P and Kir channels are about 13 e_0_, 2.2 e_0_ and 2.2 e_0_, respectively^15,16,18^.

The relatively weak voltage sensitivity of TRPV1 precluded direct measurement of q from gating current. Alternatively, the limiting-slope method^20,21^ is a classic approach to estimate q, for which voltage dependence was measured at low open probabilities. However, the voltage-driven transition of TRPV1 is allosterically coupled to activation gating^22^, with the channel open probability approaching a stable level at deep hyperpolarization (Fig. 1e and Extended Data Fig. 1) that reflected the equilibrium of activation gate between the closed and open state^23^. To better estimate q, we first calculated the derivative of the mean log open probability with respect to voltage and fitted the Boltzmann smoothing function of voltage dependence of mean log open probability (see Methods for details) (Fig. 1f), as was previously performed on the allosteric large conductance calcium activated potassium (BK) channels^23^. This approach yielded an estimated q of 0.93 e_0_, which is 30% larger than that measured from fitting a Boltzmann function to the G-V curve (0.72 e_0_), yet much smaller than that of BK channels (2.62 e_0_)^23,24^. To estimate the potential deviation of the gating charge estimate from its true value, we simulated the voltage sensitivity of TRPV1 open probability based on a simple allosteric scheme (Extended Data Figure 2a). With the hypothetical gating charge value set to span a wide range from 1.0 e_0_ to 13.0 e_0_, our estimations (Extended Data Figure 2b to 2e, dotted red curves) were close to the true value, with deviations ranging from 3.3% to 5.3% (Extended Data Figure 2f). When the method was applied to BK channels, for which total gating charge could be directly measured from the gating current, an underestimation of 7.7% was seen^23,24^.

## TRPV1 S4 does not serve as a voltage sensor

The S1-to-S4 domain of TRPV1 channels, like its counterpart in Kv channels^25^, forms a compact domain surrounding the channel pore^26,27^. In Kv channels, this domain serves as the voltage sensor^28^. To test whether TRPV1 also relies on the S1-to-S4 domain to sense V_m_, we first compared the S4 sequences (Fig. 2a). The Kv channel S4 segments contain seven regularly spaced charged residues. The corresponding residues in TRPV1 are all uncharged except for R558, which matches to R6 of Kv at the end of S4 thought to be insignificant for voltage-sensing^28^. To test for a potential contribution of R558 to voltage sensitivity, we mutated this residue to leucine. Charge-neutralization did not abolish voltage activation (Fig. 2b and c). While the G-V curve of R558L was left-shifted (Fig. 2d), the estimated q value, at 0.88 e_0_, was not significantly reduced compared to that of the wild-type channels (Fig. 2e). Similar observations were previously reported when the equivalent residue in rTRPV1 was mutated to A, L, and K^29^. Therefore, R558 in TRPV1 S4 cannot serve as the main gating charge carrier. These results showed that the S4 voltage-sensing mechanism of Kv channels is not applicable to TRPV1.

**Figure 2.**
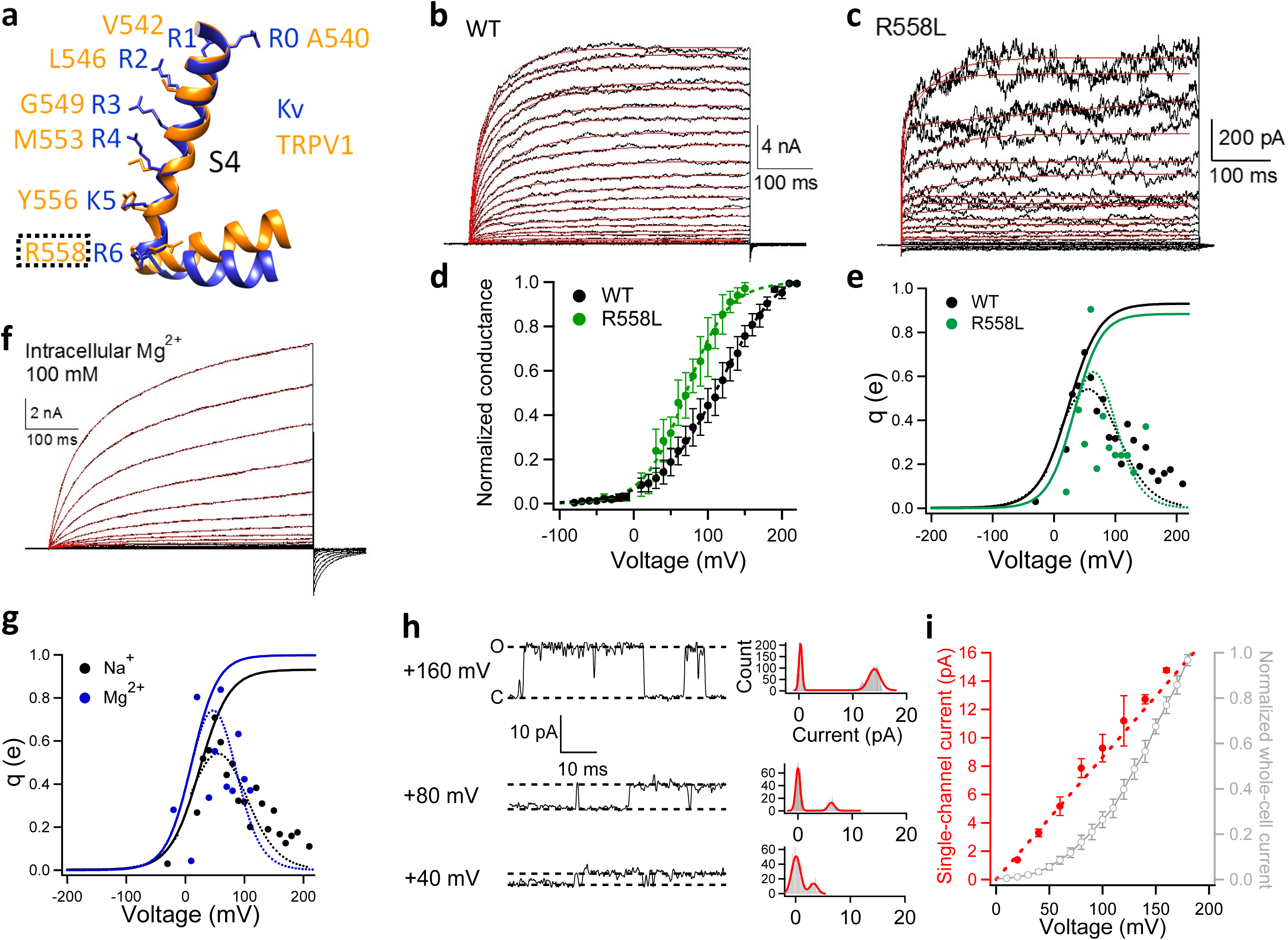
None of the three classic voltage-sensing mechanisms appears applicable to TRPV1. (**a**) Structural alignment of the S4 helixes in TRPV1 (Orange; PDB ID: 5IRZ) and Kv 1.2-2.1 chimera channel (Blue; PDB ID: 2R9R). (**b** and **c**) Representative whole-cell recordings of wild-type (WT) TRPV1 and its mutant R558L activated by depolarization with 10-mV increments from −80 mV to +220 mV and from −40 mV to +150 mV, respectively. The voltage activation time courses are well fitted to a double-exponential function (red traces). (**d** and **e**) G-V curves of WT (black) and R558L (green, V_1/2_ = 71.4 ± 12.3 mV, q_app_ = 0.96 ± 0.06 e_0_, n = 3.) in linear plot. (**e**) The voltage dependence of the derivative of mean log open probability of WT (black) and R558L (green). Dashed curve is a Boltzmann smoothing function determined from fits to mean log open probability. Solid curve is a fit of the smoothing function with a Boltzmann function at voltages where the open probability is low. (**f**) A representative whole-cell recording with Mg^2+^ as permeating ions. The voltage activation time courses are well fitted to a double-exponential function (red traces). (**g**) The voltage dependence of the derivative of mean log open probability of WT measured with intracellular sodium ions (black) and magnesium ions (blue). Dashed curve is a Boltzmann smoothing function determined from fits to mean log open probability. Solid curve is a fit of the smoothing function with a Boltzmann function at voltages where the open probability is low. (**h**) Representative single-channel recordings from an inside-out patch clamped at +40 mV, +80 mV and +160 mV, respectively. To determine the single-channel current amplitude, their corresponding all-point histograms are fitted to a double-Gaussian function (curves in red). (**i**) Voltage dependence of single-channel current amplitude (red) and macroscopic current recorded in the whole-cell configuration (gray) (n = 3-5). All statistics are given as mean ± s.e.m.

## Permeant ions do not affect voltage sensitivity of TRPV1

Since an S4-based voltage-sensing mechanism is inapplicable to TRPV1, we examined other known mechanisms. To test whether TRPV1 uses permeant ions to sense voltage like K2P channels, we took advantage of TRPV1’s low ion selectivity^8^. We reasoned that, if q is originated from permeant ion movement, switching from monovalent cations to divalent cations would likely alter the measured q value, doubling it if these cations bind to the same site(s) in the pore. When the intracellular permeant ions were switched from Na^+^ to Mg^2+^, the voltage activation and deactivation kinetics were both much slower (Fig. 2f). The G-V curve was right-shifted but without any noticeable change in shape, with an estimated total gating charge of 0.99 e_0_ (Fig. 2g). Therefore, unlike in K2P channels, permeant ions do not appear to serve as the voltage sensor for TRPV1.

## TRPV1’s voltage sensitivity is not originated from permeation block

Like K2P channels, Kir channels also lack the S1-S4 voltage sensor domain. Their currents exhibit voltage-dependence because of voltage-dependent pore block by endogenous polyamines and Mg^2+^ ions at depolarized voltages^16^. Such pore block is much faster than the resolution of patch-clamp recordings so that, when it occurs, the single-channel conductance is reduced. As a result, the single-channel current amplitude exhibits similar voltage dependence as the macroscopic current^30^. To test whether TRPV1 employs this pore-block mechanism for sensing voltage, we compared single-channel current and macroscopic current recorded at voltages where TRPV1 was clearly activated by depolarization (Fig. 2h and i). It was obvious that the single-channel current amplitude was linearly dependent on voltage and did not follow the voltage dependence of macroscopic current (Fig. 2i). Moreover, in inside-out patch recordings the voltage dependence did not decrease upon continuous perfusion of the bath solution for about three minutes (Extended Data Fig. 3). In comparison, Kir channels lost the inward-rectification feature within two minutes of patch excision due to wash-off of endogenous blockers^16^. Therefore, unlike Kir channels, voltage sensitivity of TRPV1 is unlikely originated from pore block.

## TRPV1 voltage-dependent gating behavior resembles a concerted transition

As none of the known voltage-sensing mechanisms in the VGL channel superfamily is applicable to TRPV1, an unorthodox mechanism exists. We gained the first glimpse of such a mechanism from kinetic measurements. Voltage activation of Kv2.1 channel exhibited a sigmoidal time course^31^ (Fig. 3a), where there was an initial delay in macroscopic current upon depolarization (marked by red arrow). This delay is due to multiple closed states Kv channels must traverse before reaching the open state^20^ (Fig. 3c, SCHEME I). In contrast, voltage activation of TRPV1 nicely followed a double-exponential time course (Fig. 3a and Fig. 2b, red solid traces), with no detectable delay in current onset. Indeed, time courses predicted by gating schemes for Kv channel in the form of SCHEME I poorly described the voltage activation kinetics of TRPV1 (Fig. 3a, blue dash trace). In addition, it is well established that for Kv channels holding V_m_ at more hyperpolarized voltages would produce a longer delay in current activation upon depolarization. Such a longer delay, named Cole-Moore shift^32^, is again due to the existence of multiple closed states preceding the open state. When TRPV1 channels were voltage-clamped at various hyperpolarized voltages, there was no detectable Cole-Moore shift (Fig. 3b). The absence of both initial delay in macroscopic current activation and Cole-Moore shift, as well as the double-exponential current time course, can be explained by an allosteric model as shown in Figure 3c, SCHEME II, in which there is a single concerted voltage-dependent transition that may represent either the C↔O transition or a separate transition that is allosterically coupled to the C↔O transition. (The observation of a double-exponential activation time course rules out a simpler two-state C↔O model for voltage-dependent activation.) Admittedly, more complex models, for example, with multiple rapid voltage-dependent transitions and a single rate-limiting voltage-independent transition right before the first opening transition, could also produce similar observations in channel activation time course and the absence of a Cole-Moore shift^33^. However, none of the currently available experimental evidence requires such a complex gating model and, as discussed above, TRPV1 voltage activation involves an allosteric coupling. Furthermore, the simple allosteric model in Figure 3c, SCHEME II, is consistent with our results described below.

**Figure 3.**
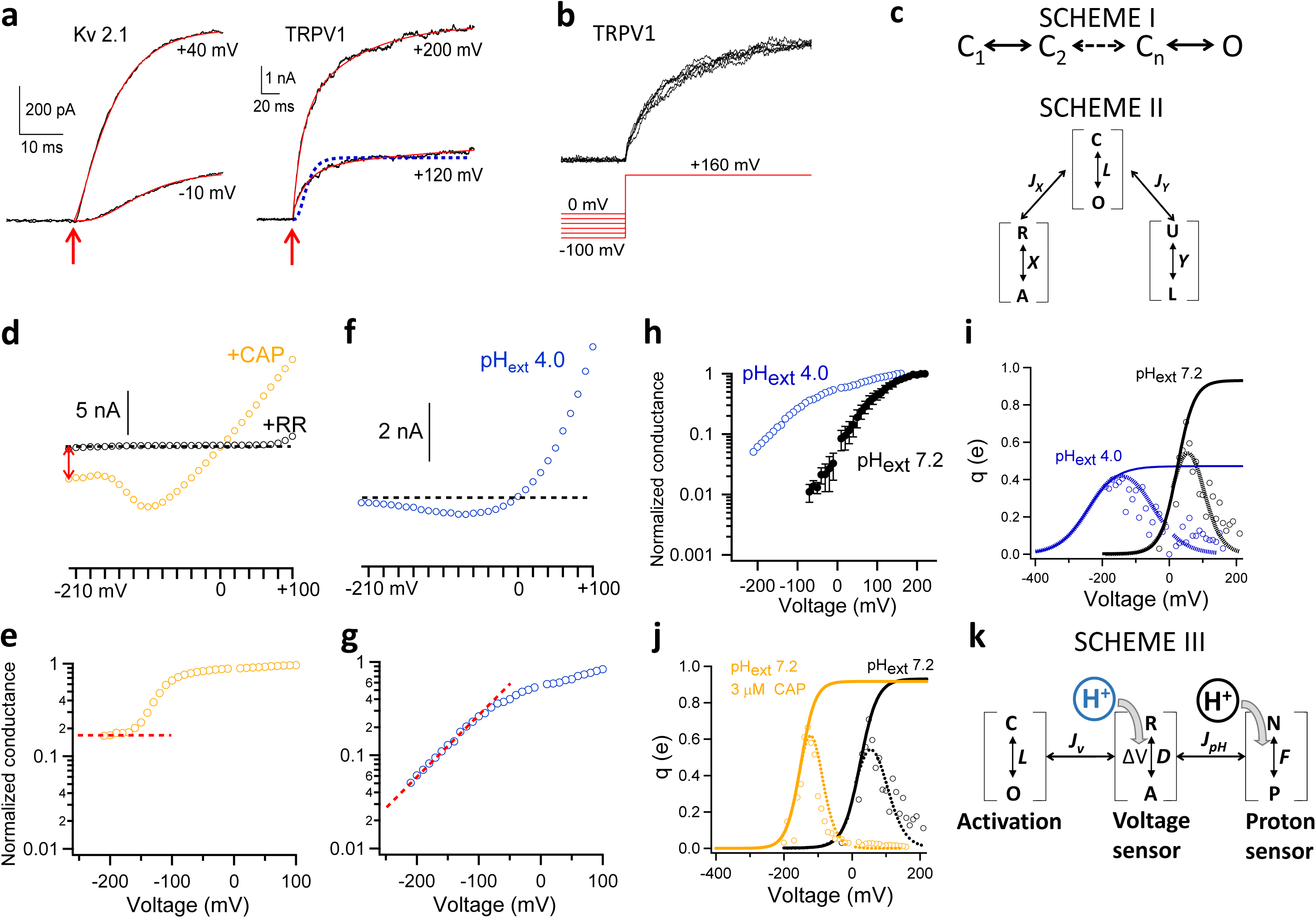
Voltage-sensing of TRPV1 involves the proton activation machinery. (**a**) Voltage activation kinetics of Kv 2.1 channels exhibit an initial delay between depolarization (marked by an arrow in red) and the onset of macroscopic current. No such a delay was observed in voltage activation of TRPV1. The activation time course of Kv 2.1 is fitted to Equation (2) (based on SCHEME I in (**c**); also see Methods in Supplementary materials), whereas a double-exponential function is used to fit that of TRPV1 (solid curves in red). SCHEME I predicted the activation time course of TRPV1 shown by the dashed trace in blue. (**b**) No Cole-Moore shift was observed in voltage activation of TRPV1. No delay in current onset can be observed from macroscopic currents recorded at +160 mV after a 2-s prepulse from 0 mV to −100 mV with 20-mV increments. (**c**) Gating SCHEME I and II for voltage activation of Kv channels and TRPV1, respectively. (**d**) A representative recording of the voltage dependence of TRPV1 in whole-cell configuration in the presence of 10 µM CAP. Orange circles represent the steady-state current amplitude at the corresponding voltages. Leak current was measured from the same patch in the presence of ruthenium red (20 µM) to block TRPV1 (black circles). Current through TRPV1 at - 210 mV is indicated by a double-arrow in red. A dashed line in black indicates the zero-current level. (**e**) Voltage dependence of the normalized conductance calculated from the recording in (**d**). A dashed line in red indicates a steady-state level was reached at deeply hyperpolarized voltages beyond −160 mV. (**f**) A representative recording of the voltage dependence of TRPV1 in whole-cell configuration in the presence of extracellular pH 4.0. (**g**) Voltage dependence of the normalized conductance calculated from the recording in (**f**). A dashed line in red indicates the conductance level kept decreasing at deeper hyperpolarization. (**h**) The semi-logarithmic plot for the voltage dependence of normalized mean conductance measured from steady-state current in whole-cell recordings at extracellular pH 4.0 and pH 7.2, respectively. (**i**) The fits to the derivative of the mean log open probability at extracellular pH 4.0 and pH 7.2, respectively. (**j**) The fits to the derivative of the mean log open probability with and without 3 µM capsaicin, respectively. (**k**) A gating scheme where the voltage- and H^+^-dependent transition couples allosterically to channel opening.

## Identifying the voltage-dependent transition

Besides membrane depolarization, TRPV1 can be allosterically activated by distinct physical and chemical stimuli, many of which have been better characterized both structurally and functionally^22,26,27,34^. A multi-allosteric gating model initially proposed for BK channels^35,36^ can adequately describe TRPV1 activation^37-40^. The polymodal nature of TRPV1 activation offered an opportunity to identify the voltage-dependent transition. For a general allosteric system, there exist three possible scenarios as shown by SCHEME II (Fig. 3c) that are experimentally distinguishable by measuring open probability, P_o_, at deeply hyperpolarized voltages:

A. Voltage directly controls the C↔O transition. In this scenario, the channel can be tightly held in the C state by hyperpolarization; no other stimuli should be able to evade voltage’s control over the C↔O transition to cause channel activation.
B. Voltage controls a transition, R↔A, that is allosterically coupled to the C↔O transition, and another stimulus, *X*, also activates the channel through the same transition. In this scenario, hyperpolarization can effectively prevent channel activation by *X* but not by other stimuli (for example, *Y*).
C. Voltage controls a distinct transition, independent of other stimuli, that is allosterically coupled to the C↔O transition. In this scenario, all other stimuli can efficiently activate the channel at hyperpolarized voltages.

To distinguish these scenarios, we measured NP_o_ from single-channel events at deep hyperpolarization down to −210 mV in the absence or presence of another stimulus. We found that, in the absence of another stimulus, NP_o_ appeared to approach a steady level at deeply negative voltages (Extended Data Fig. 1). In agreement with previous reports^37,38,40,41^, the observed baseline P_o_ level was very low, with an upper limit estimate near 0.01 at room temperature (Fig. 1e). Nonetheless, the presence of a voltage-independent baseline P_o_ is inconsistent with Scenario A, suggesting that the C↔O transition itself must be associated with little charge movement and have an intrinsic open probability (P_o_intrinsic_) of less than 0.01.

In the presence of a saturating concentration (3 µM) of capsaicin, TRPV1 current was readily detectable even at −210 mV (Fig. 3d, red arrow). The current could be almost completely blocked by ruthenium red (Fig. 3d, black circles), confirming that contamination by leak current must be small. The estimated P_o_ plateaued above 0.1 when voltage was hyperpolarized below −160 mV (Fig. 3e, red dash line; see also Fig. 1c). Similar to previous reports^40,41^, the presence of a stable P_o_ substantially above P_o_intrinsic_ indicated that capsaicin could allosterically promote the C↔O transition even at deep hyperpolarization. Therefore, voltage activation cannot be carried out through the same transition promoted by capsaicin, i.e., if Scenario B is true, capsaicin cannot be stimulus *X*. Indeed, as capsaicin induces TRPV1 activation by linking S4 to the S4-S5 linker^42-44^, these results offered further supports for the conclusion that voltage activation does not involve S4. Furthermore, the observation of a voltage-independent, elevated P_o_ in the presence of capsaicin also validated the conclusion that voltage cannot work through the C↔O transition to open the channel, i.e., Scenario A is invalid.

Like capsaicin, we found that many other TRPV1 activation stimuli, including heat^45^, were also effective at deeply hyperpolarized voltages (see, for example, Fig. 1c). However, the situation was different when extracellular protons were used as the activator. We found that even in the presence of a saturating concentration of protons (at pH 4.0), current decreased towards zero at hyperpolarizing voltages (Fig. 3f). No plateau could be observed in the P_o_–V plot (Fig. 3g); instead, P_o_ kept decreasing towards P_o_intrinsic_ (Fig. 3h). It is known that protons partially block TRPV1 permeation, which also exhibits a shallow voltage dependence^46^. However, proton blockade had been corrected for before calculating P_o_ (Extended Data Fig 4.). Therefore, these observations fit the prediction of Scenario B in that deep hyperpolarization can shut down proton activation, identifying that voltage works though the proton activation pathway. These results also voided Scenario C by identifying protons as the stimulus *X*.

## Location of the gating charges

While unanticipated, the finding that voltage sensitivity of TRPV1 resides in the proton activation pathway might not be surprising. It is well established that proton activation involves protonation of charged residues in the outer pore^47^, a region intimately involved in TRPV1 activation gating and known to undergo substantial conformational rearrangements^48^. The outer pore is also noticeably rich in charged residues (Extended Data Fig. 5a). If some of these charged residues locate within or near the V_m_ field, their movements during channel activation will impart voltage sensitivity to the process^5^. Indeed, when we simultaneously neutralized titratable negative charges by changing the extracellular pH to 4.0, the total gating charge estimate decreased to 0.47 e_0_ (Fig. 3i). Given that the pKa value for the sidechain of negatively charged Glu or Asp is around 4, at extracellular pH 4.0 only about half of these charged residues would be neutralized. Therefore, we reasoned that though we observed only an around 50% reduction of q at acidic pH, nearly all of the 0.93 e_0_ total gating charge could be collectively contributed by titratable acidic residues. Consistent with a previous report^49^, no obvious change in the total charge movement could be confidently identified from channels carrying single or double charge neutralization mutations (Extended Data Fig. 5b and 5c), likely because their individual impacts to q were too small to stand out of the noise. These mutagenesis tests confirmed that the total gating charge of about 1 e_0_ is not carried by any single charged residue (which would require the residue to move nearly 1/4 of the whole V_m_ field during activation). Instead, it is most likely that multiple charged residues move collectively during voltage gating. As a negative control we observed no change in total gating charge in the presence of the classic TRPV1 agonist capsaicin (Fig. 3j).

Based on these findings, we tentatively placed the titratable total gating charge at the voltage-driven R↔A transition that is also affected by protons (SCHEME III, Fig. 3k). A gating model incorporating this feature predicted behaviors that qualitatively resemble our experimental observations (Extended Data Fig. 6).

## Structural mechanism for voltage/proton activation

The TRPV1 closed-state structure and vanilloids/toxin-activated structures have been resolved by cryo-EM^26,27,34^. However, these structures are missing a part of the outer pore important for voltage and proton activation; in addition, no voltage/proton-induced open structure is currently available. To understand the structural basis for voltage/proton-dependent gating, we first modeled the proton-induced open state structure using Rosetta structural modeling suite (see Methods and Extended Data Fig. 7). Comparison between the proton-activated state model and the closed state structure suggested two major conformational changes. Firstly, the selectivity filter region moved away from the central axis to yield an open conformation of this upper gate (Fig. 4a, gray to red), similar to DkTx-induced conformational changes observed in the cryo-EM studies^27,34^. Secondly, the pore turret regions moved substantially closer to each other upon protonation (Fig. 4a, dark gray to cyan), which has been previously observed by fluorescence recordings during heat activation^50^. Associated with these structural changes are relocation of multiple charged residues.

**Figure 4.**
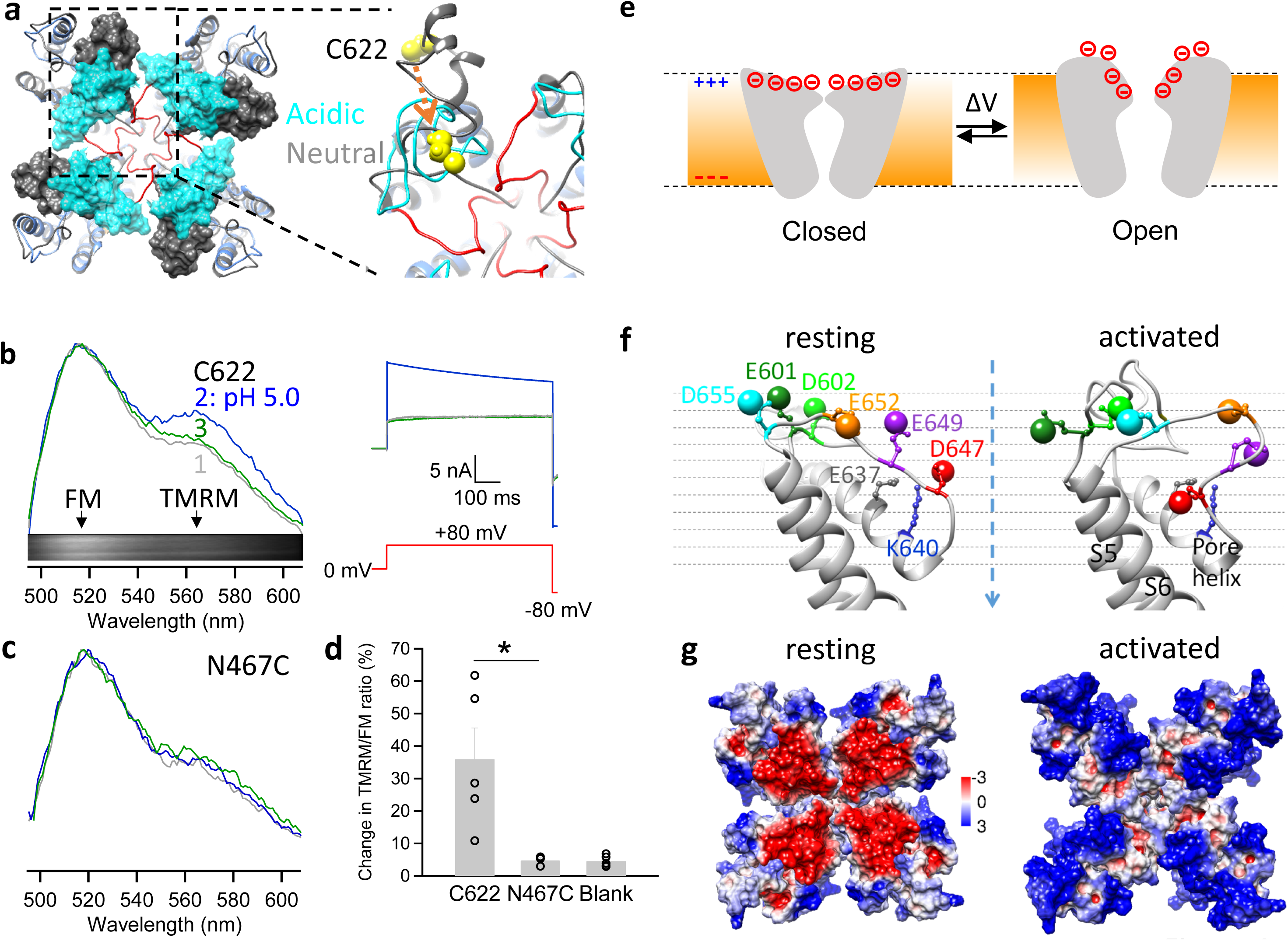
Structural mechanism underlying voltage sensitivity in TRPV1. (**a**) The pore region of TRPV1 modeled under acidic (pH 4.0) and neutral conditions based on the cryo-EM structure in the closed state (PDB ID: 3J5P). For the model under acidic pH, its selectivity filter and the linker to S6 are colored in red. The turrets under acidic and neutral pH are shown in cyan and dim gray, respectively. Atoms of C622 on the turret are shown as spheres in yellow. An arrow in orange indicates the direction of C622 movement upon acidification. (**b**) Proton-induced conformational changes at the turrets measured by whole-cell patch fluorometry from FM and TMRM attached to C622 sites. 1, 2 and 3 indicate the emission spectra and their corresponding current recordings at neutral pH, perfusion of a pH 5.0 solution and wash off, respectively. The emission spectra were normalized to the emission peak of FM. Inset shows a representative spectral image of fluorescence emission. (**c**) Similar measurements as (**b**) at the N467C site in the S1-S2 linker as a negative control. (**d**) Percentage change in the TMRM/FM intensity ratio measured from fluorophores labeled at C622, N467C and in blank cells expressing no TRPV1 channel, respectively. *, P < 0.05. n = 3-to-8. All statistics are given as mean ± s.e.m. (**e**) Left, a schematic diagram illustrating depolarization-induced collective movements of charged residues and dipoles in the outer pore region. Right, comparison of TRPV1 structure in the closed state (grey) and the activated state model (blue). Helices are presented as pipes. Conformational changes in charged residues and helixes underlie the unorthodox voltage-sensing mechanism in TRPV1. (**f**) Locations of outer pore charged residues in the structurally aligned resting and activated state models. The locations of negative charges are approximated by the oxygen atoms in the sidechain (shown as a large sphere). The dashed arrow in blue indicates the central axis through the pore (also defined as the z axis of the models). (**g**) The surface electrostatic potential of the resting and activated states. Positive and negative potentials (in kT/e) are colored in blue and red, respectively.

To functionally verify the predicted outer pore movement, we employed FRET-based patch fluorometry to simultaneously record conformational and functional states of TRPV1 channels^50,51^. Our structural models predicted that the distance between C622 residues of neighboring subunits (measured at C_β_ atoms; Fig. 4a, yellow) is reduced by about 13 Å in proton-induced activation. This large distance change would be readily detectable using fluorescein maleimide (FM) and tetramethylrhodamine maleimide (TMRM) as a FRET pair^50^. Extracellular acidification (pH 5.0) elicited a large current from the fluorophore-labeled channels (Fig. 4b). Fluorescence imaging from the same channel population revealed an increase in the TMRM/FM intensity ratio that indicated an increase in FRET efficiency (Fig. 4b). This change in emission spectrum was absent when fluorophores were attached to cysteine introduced at N467 on the S1-S2 linker, or at non-specific sites on untransfected cells (Fig. 4c and d). Therefore, results from the FRET experiments supported the prediction that the turrets move toward each other in the proton-activated state.

How does the observed outer pore conformational changes produce voltage-sensitive gating? While much of the detail remains to be elucidated, it is conceivable that relocation of some negatively charged residues within the V_m_ field might be the origin of the gating charge (Fig. 4e). Indeed, from the activated-state model, we could identify multiple negatively charged residues in the outer pore region that exhibited complex and collective movements (Fig. 4f): D602, D647, E652 and D655 moved downward along the z axis, E601 and E649 moved upward (Extended Data Table 2). Based on the simplified assumption of a uniform V_m_ field that spans a 30 Å vertical depth, these charge movements alone would be more than sufficient to produce 1 e_0_. It was also noticed that the modelled gating rearrangements would lead to changes in the electrostatic potential surface of the outer pore, in which a strong negative potential in the resting state dissipates as the channel transitions into the activated state (Fig. 4g). Such conformational changes present a plausible picture of the voltage-sensitive gating process: it is the collective gating rearrangements of negatively charged residues and the local electric field that produce an unorthodox mechanism of voltage-sensing in TRPV1.

## Discussion

Our study suggested that voltage-sensing in TRPV1 originates from conformational changes of the outer pore, where titratable acidic residues collectively contribute to the total charge movement. Because that the pKa value for the sidechain of negatively charged Glu or Asp is around 4, extracellular acidification to a pH level of 4.0 would only neutralize about half of the charges in these charged residues. It was experimentally infeasible to further decrease extracellular pH in patch-clamp recordings to fully neutralize all the negative charges, though it is most likely that the about 1 e_0_ total gating charge is collectively contributed by titratable acidic residues. Upon depolarization, these residues appear to undergo complex rearrangements that are directly coupled to opening of the activation gate, most likely the nearby upper gate at the selectivity filter^26,27,34^ (Fig. 4g). The pore turret is another outer pore channel structure seen to move substantially in our model as well as site-directed fluorescence measurements in our previous studies^50,52^ and the present study (Fig. 4b-d). Details on the nature of pore turret movements remain unclear, as this segment is missing from the available TRPV1 cryo-EM structures; nonetheless, the equivalent part of the orthologous TRPV2 is seen to move substantially between the closed and open states^53^.

The TRPV1 outer pore is a known hot spot for gating modulation of this polymodal receptor. Small cations such as proton, Na^+^ and Mg^2+^/Ba^2+^/Ni^2+^/Gd^3+^, as well as large peptide toxins such as DkTx, RhTx and BmP01 all bind to this region to exert their strong gating effects^48^. Capsaicin activation has been recently found to affect the outer pore^44^. Heat also induces large conformational changes of the outer pore^50^. Exploiting charged residues and dipole movements in this gating structure would allow V_m_ to function essentially as a “master gain setter” to tune the channel’s sensitivity to all major stimuli (see Fig. 1c), a feature that could be of high practical significance for this nociceptor. For example, under pathological conditions such as bone cancer, TRPV1-expressing dorsal root ganglion neurons exhibit higher excitability^54^. The resting membrane potential of these neurons is depolarized, which leads to higher nociceptive activities of TRPV1, contributing to the severe and intractable pain in the bone cancer. When TRPV1 activity is modulated by hyaluronan, firing frequency of DRG neurons is also changed^55^. In this regard, understanding the voltage-sensing mechanism may facilitate future pharmaceutical efforts targeting this pain sensor.

Comparing to many noxious stimuli for TRPV1 such as capsaicin and heat, voltage is a relatively mild activator. The weak voltage sensitivity ensures that TRPV1 does not act as a dedicated V_m_ sensor, which is critical for its nociceptive functions. The total charge movement of about 1 e_0_, though only a fraction of that in classic S4-based voltage-gated channels, can effectively drive man-made transistors in conventional electronic devices^4^ and many biological functions of membrane proteins. As pointed out by Hodgkin^56^, gating charge movement absorbs energy from electrical signaling and acts as a load of the system, therefore evolution tends to optimize its utility. We showed recently that weak voltage dependence is a crucial feature that allows TRPV1 to respond to diverse noxious stimuli including heat, whereas the high voltage sensitivity of Kv channels effectively prohibits the channel’s intrinsic heat sensitivity^45^. It is plausible that similar trade-offs can be found in other voltage-regulated polymodal TRP channels.

When the S4 of Kv channels is functionally decoupled from the pore by introducing mutations, a weakly voltage-dependent (q of ∼1.8 e_0_), concerted transition is revealed^57^. In CNG channels, S4 (containing multiple charged residues) appears to be naturally decoupled from the gate^58^, and the channel’s intrinsic weak voltage dependence (q of ∼ 0.74 e_0_) likely comes from outer pore movement^59,60^. Therefore, it is likely that the outer pore-mediated voltage-sensing mechanism is broadly used in ion channels. Indeed, many TRP channels exhibit similar shallow voltage dependence, sparse charged residues in S4 but richly distributed charged residues in the outer pore, a region seen to under tremendous evolutional selection^61^. While activation of these channels is also critically regulated by V_m_, their voltage-sensing mechanism is unknown (Fig. 1d, section in red). The unorthodox voltage-sensing mechanism we identify in TRPV1 may help elucidate the function of these channels and other membrane proteins.

## AUTHOR CONTRIBUTIONS

F.Y., X.L.Z., B.H.L. and X.X. conducted the experiments including patch-clamp recording, mutagenesis, molecular modeling and data analysis; V.Y.-Y. supervised molecular docking and revised the manuscript; J.Z. and F.Y. prepared the manuscript; J.Z., V.Y.-Y. and F.Y. conceived and supervised the project, participated in data analysis and manuscript writing.

## ACKNOWLEDGEMENTS

We are grateful to Fred Sigworth for reading the manuscript and providing advice, and to our lab members for assistance and discussion. This work was supported by funding from National Institutes of Health (R01NS072377 and R01NS103954) to J.Z. and V.Y.Y., American Heart Association (14POST19820027) to F.Y. and American Heart Association (16PRE26960016) to B.H.L. This work was supported by National Science Foundation of China (31741067 and 31800990) to F.Y. This work was also supported by the Core Facilities in Zhejiang University School of Medicine, including the Olympus FV3000 fluorescence imaging microscope and the bioinformatics computation platform.

## DATA AVAILABILITY

All data needed to evaluate the conclusions in the paper are present in the paper and/or the Supplementary Materials. Additional data available from authors upon request.

## MATERIALS AND METHODS

### Molecular biology and cell transfection

cDNA of Murine TRPV1 (a gift from Dr. Michael X. Zhu, University of Texas Health Science Center at Houston) and Kv2.1 (a gift from Dr. Jon Sack, University of California, Davis) were used in this study. To facilitate identification of channel-expressing cells, eYFP and GFP were fused to the C-terminus of TRPV1 and the N-terminus of Kv2.1, respectively. Tagging of these fluorescence proteins did not alter channel functions, as previously reported^1,2^. QuickChange II mutagenesis kit (Agilent Technologies) was used to generate point mutations. All mutations were confirmed by sequencing.

HEK293T cells, purchased from and authenticated by American Type Culture Collection (ATCC), were cultured at 37°C with 5% CO_2_ in a Dulbecco’s modified eagle medium with 10% fetal bovine serum and 20 mM L-glutamine. Transient transfection with cDNA constructs was done with Lipofectamine 2000 (Life technologies) following manufacturer’s protocol. Patch-clamp recordings were performed 1-2 days after transfection.

### Chemicals

All chemicals such as capsaicin were purchased from Sigma-Aldrich.

### Electrophysiology

Patch-clamp recordings were performed with a HEKA EPC10 amplifier with PatchMaster software (HEKA) in inside-out or whole-cell configuration. Patch pipettes were prepared from borosilicate glass and fire-polished to resistance of ∼4 MΩ. For whole-cell recording, serial resistance was compensated by 60%. A normal solution containing 130 mM NaCl, 10 mM glucose, 0.2 mM EDTA and 3 mM HEPES (pH 7.2) was used in both bath and pipette. For solutions with 6.0 < pH < 7.2, HEPES was used as the pH buffer. For solutions with pH ≤ 6.0, HEPES was replaced by 3 mM 2-(N-morpholino)ethanesulfonic acid (MES). When intracellular Na^+^ ions were replaced by Mg^2+^ ions, there was no Mg^2+^ in the extracellular solution. To determine a G-V curve, the membrane potential was first clamped at −80 mV for 10 ms and then switched to another clamping voltage stepping from −210 mV to +210 mV with a 10-mV interval for 500 ms, which was then switched to −80 mV. Current amplitude at the steady state during the last 100 ms of voltage steps was averaged to construct the G-V curve. Current signal was filtered at 2.9 kHz and sampled at 10 kHz.

To apply solutions containing capsaicin or other reagents during patch-clamp recording, a rapid solution changer with a gravity-driven perfusion system was used (RSC-200, Bio-Logic). Each solution was delivered through a separate tube so that there was no mixing of solutions. Pipette tip with a membrane patch was placed directly in front of the perfusion outlet during recording. Each membrane patch was recorded for only once.

### Temperature control and monitoring

The bath solution was heated using an SHM-828 eight-line heater driven by a CL-100 temperature controller (Harvard Apparatus). We placed a TA-29 miniature bead thermistor (Harvard Apparatus) about 1 mm from the pipette tip to monitor local temperature change. Temperature readout from the thermistor was fed into an analog input port of the HEKA patch-clamp amplifier and recorded simultaneously with channel current. The speed of temperature change was set at a moderate rate of about 0.3 °C/s to ensure that heat activation reached equilibrium during the course of temperature change and the current was recorded at steady state. When the experimental temperature was not controlled, recordings were conducted at room temperature at 24°C. Temperature variation was less than 1 °C as monitored by a thermometer.

### Site-directed fluorescence recordings

Structural changes in the turret and other extracellular regions were monitored with site-directed fluorescence recordings as previously described ^3,4^. Briefly, fluorescein maleimide (FM) and tetramethylrhodamine maleimide (TMRM) were used to irreversibly label a native cysteine C622 in the turret region (after removing another native cysteine by mutation C617A), or an introduced cysteine N467C in the S1-S2 linker region after removing both C617 and C622 with an alanine mutation. Fluorescence resonance energy transfer (FRET) between FM and TMRM was measured from voltage-clamped HEK293 cells imaged with an inverted fluorescence microscope (Nikon TE2000-U) using a 40X oil-immersion objective (NA 1.3). An argon laser (Spectra-Physics) was used to provide the excitation light, with the exposure time controlled by a Uniblitz shutter synchronized with the camera by the PatchMaster software through HEKA amplifier. Spectral measurements were performed with an Acton SpectraPro 2150i spectrograph in conjunction with a Roper Cascade 128B CCD or an Evolve 512 EMCCD camera. The rates of photobleaching FM and TMRM by the excitation light were quantified separately from fluorophores attached to the channel and used to correct for photobleaching during FRET experiments. Proton-dependent changes in fluorescence intensity were also measured and corrected. FRET was quantified from the enhancement of acceptor fluorescence emission due to energy transfer^1,5^. FRET measurements were done with cells that were patch-clamped throughout the recording. Proton-induced current potentiation and FRET change were recorded simultaneously.

### Molecular Modeling

To model the pore region of TRPV1 under neutral and acidic extracellular pH, membrane-symmetry-loop modeling was performed using the Rosetta molecular modeling suite6. First, to model the pore region under neutral pH, coordinates of S1-S5, Pore helix, and S6, were taken from TRPV1 channel structures (PDB ID: 3J5P) and kept rigid during loop modeling and finally relaxed. The loop regions between S1 and S2, S2 and S3, S5 and pore helix (turret), the selectivity filter and its linker to S6 were modeled *de novo* in each subunit of the tetramer. During the first round of modeling, Rosetta’s cyclic coordinate descent (CCD) loop relax protocol^7^ was used and the top 20 cluster center models were passed to the second round. During the second to third rounds of modeling, kinematic (KIC) loop relax protocol^7,8^ was used and the top 20 cluster center models were passed to the next round. During the rest rounds of modeling, kinematic loop relax protocol was used and the top 20 models by score were passed to the next round. From 10,000 to 20,000 models were generated in each round. Models reported here represent the lowest energy models from the last round of iterative loop relax. The TRPV1 models we generated clearly entered an energy funnel through the ten rounds of loop modeling (Extended Data Figure 6), suggesting that our final model is located at the bottom of energy well. Top 10 models with lowest energy after the last round of loop modeling exhibited good structural convergence (Extended Data Figure 6).

To model the pore region under acidic pH, we followed the established protocols in Rosetta^9,10^. Briefly, we modified the membrane energy function terms^11,12^ as described in these reports. Then we added the following command during the loop modeling:

~~~
-pH_mode true
~~~

We set the pH value to be 4 by adding this command:

~~~
-value_pH 4
~~~

Rounds of CCD and KIC loop modeling were performed with these commands to obtain the model of TRPV1 pore region under acidic pH.

The quality of models under neutral and acidic pH were rigorously assessed by four independent protein structure analysis and verification methods^13-16^. The metrics of our models were similar to those derived from cryo-EM and crystallography (Extended Data Table 1).

All the molecular graphics of TRPV1 models were rendered by UCSF Chimera^17^ software version 1.11.

## Data analysis

To characterize the steady-state G-V curves, a single-Boltzmann function was used:

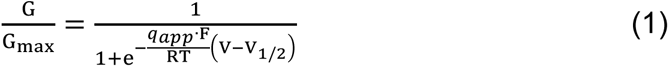

where G/Gmax is the normalized conductance, V_1/2_ is the half-activation voltage, q_app_ is the apparent gating charge and F is Faraday’s constant, R is gas constant, and T is temperature in Kelvin.

To describe the voltage activation kinetics of Kv 2.1 channels, the following equation was used:

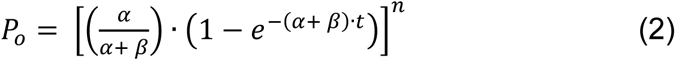

where α and β are microscopic forward and backward transition rates, respectively, n is the number of independent transitions the channel must transverse before opening. For Kv 2.1 channels we set n to be 4.

To estimate the total gating charge from the voltage dependence of channel activation, we followed the method employed in the study of BK channels^18^. Briefly, the mean activation charge displacement q_a_ was first calculated by

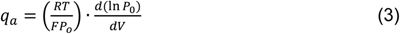

A smoothing function in the form of equation (1) was imposed on the voltage dependence of Ln(Po), and then the total gating charge was estimated by fitting the foot of the q_a_-V curve with a function:

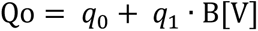

where B[V] is a Boltzmann function in the form of equation (1), q_0_ and q_1_ represent the amplitudes of the voltage-independent and voltage-dependent components of Q_o_ (total gating charge), respectively.

Open probabilities of TRPV1 channel lower than 1% were often difficult to quantify reliably; for this reason, we analyzed voltage dependence at lower open probabilities using NPo from single-channel recordings.

### Multi-allosteric gating model

The approach we used for the study of TRPV1 polymodal gating was inspired by the work on Ca^2+^-sensitive voltage-gated BK channels by Aldrich, Horrigan, and Cui ^18-20^. Similar to these earlier studies, our approach to a general gating model for TRPV1 assumed that the closed-to-open transition, C←→O, and the transition induced by a given activation stimulus, R←→A, are both reversible transitions that are coupled allosterically. The details in our modeling have been described previously^21^.

### Statistics

All experiments have been independently repeated for at least three times. All statistical data are given as mean ± s.e.m.. Two-tailed Student’s *t*-test was applied to examine the statistical significance. N.S. indicates no significance. ***, *p* < 0.001.

## EXTENDED DATA FIGURE LEGENDS

**Extended Data Figure 1.**
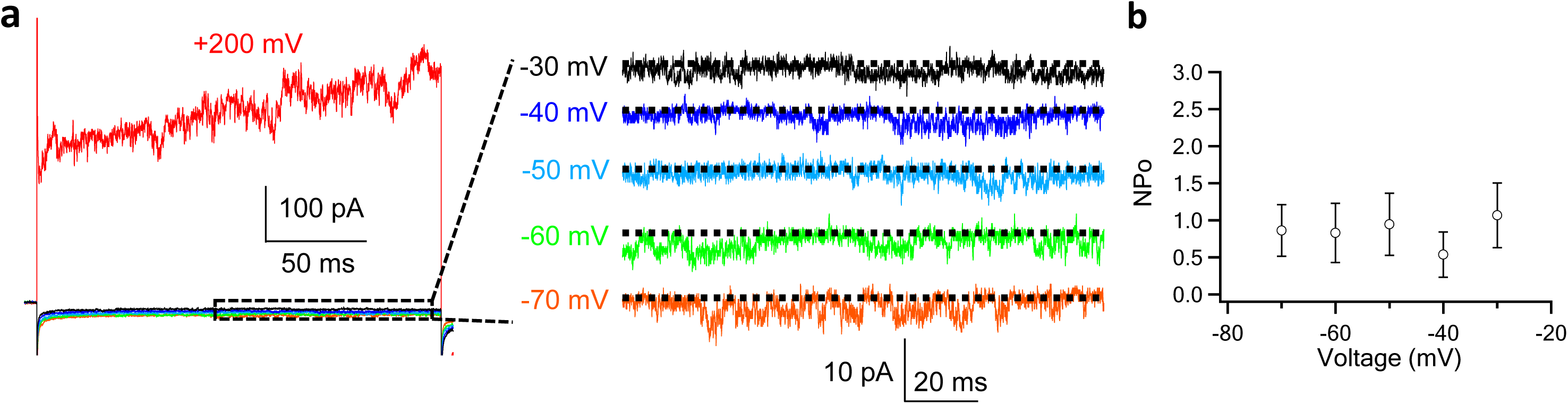
Patch-clamp recording of TRPV1 at hyperpolarized voltages. (a) A representative inside-out patch recording with voltage stepped from −70 mV to +200 mV. Single-channel activities were clearly discernable at hyperpolarized voltages. The open probability did not further decrease with deeper hyperpolarization beyond −30 mV. (b) Voltage dependence of NPo beyond −30 mV remained at a stable level (n = 4).

**Extended Data Figure 2.**
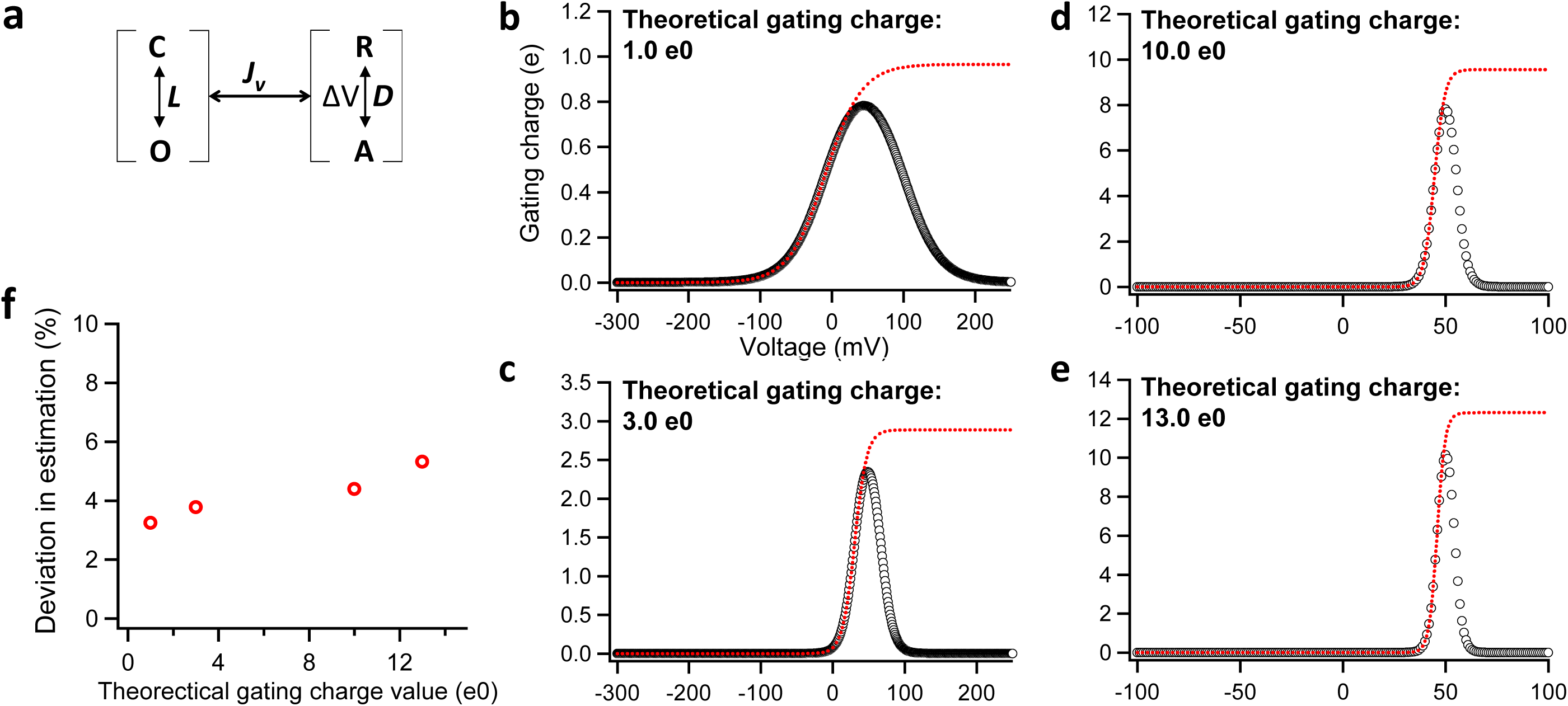
Estimation of the deviation between the true value of total gating charge and the measured values. (a) The simple allosteric voltage gating scheme employed for the estimation. (b to e) The voltage dependence of open probability was simulated with the gating scheme in (a) and the theoretical total gating charge was set to 1.0 e_0_, 3.0 e_0_, 10.0 e_0_ and 13.0 e_0_, respectively. The simulated voltage dependence of open probability (open circles in black) was further analyzed as described in Method and in Figure 1 (f) to derive the voltage dependence of total gating charge (dotted curve in red). (f) The measured total gating charge (the maximum value of the dotted curve in red) was compared to the theoretical true values to calculate the deviation in total gating charge estimation.

**Extended Data Figure 3.**
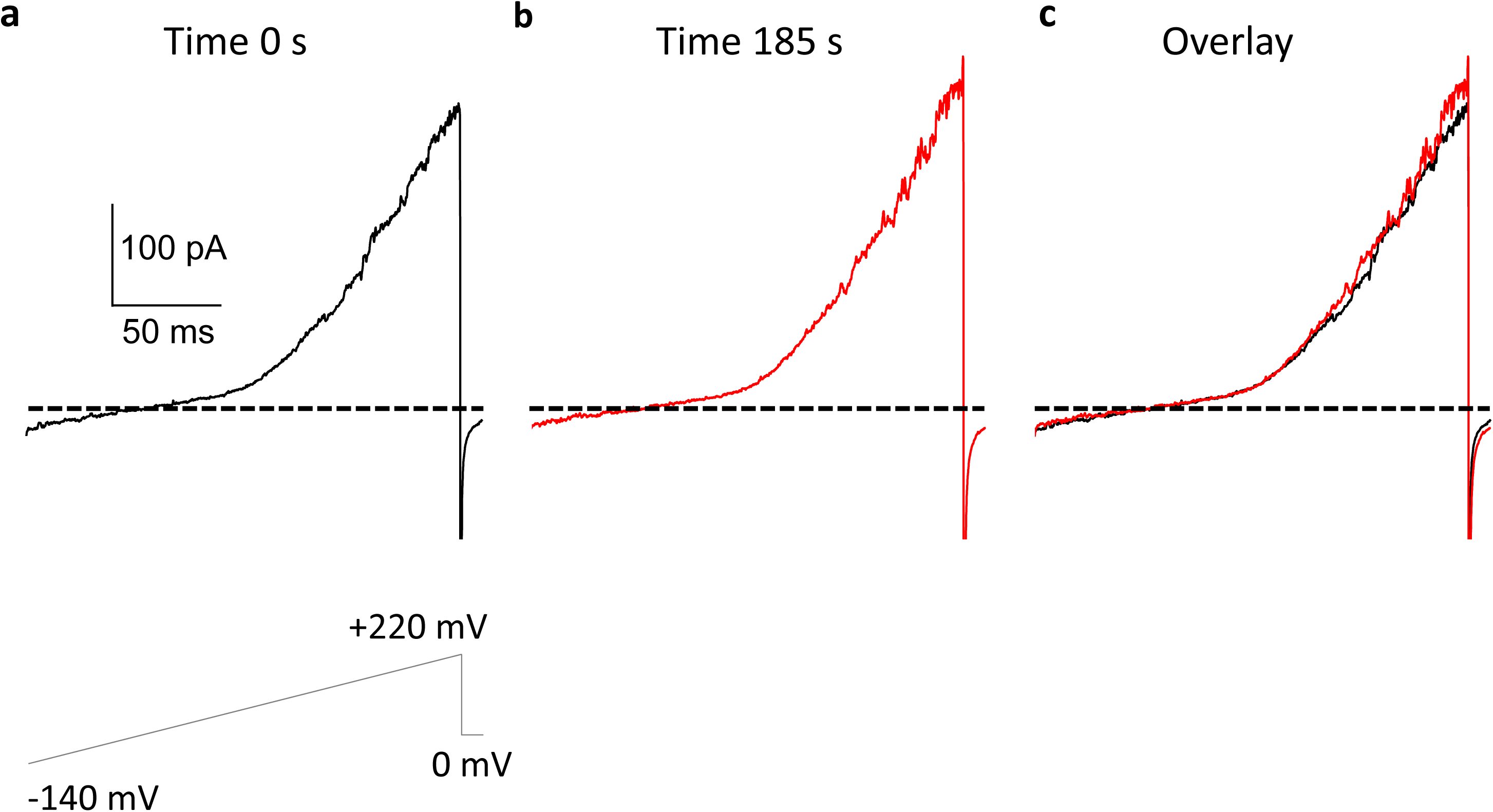
The voltage activation of TRPV1 does not diminish upon washing off intracellular factors. (**a**) A representative inside-out patch recording of TRPV1 activated by depolarization during a voltage ramp. This patch was under constant perfusion of bath solution on its exposed intracellular side. (**b**) After constant wash-off for more than three minutes, the patch exhibited a similar current response to the same voltage ramp as in (**a**). (**c**) The overlay of voltage-activated current traces at time 0 s and 185 s demonstrates that washing off intracellular factors did not alter voltage sensitivity in TRPV1. All experiments were performed at 22°C. Only transmembrane voltage was used to activate TRPV1.

**Extended Data Figure 4.**
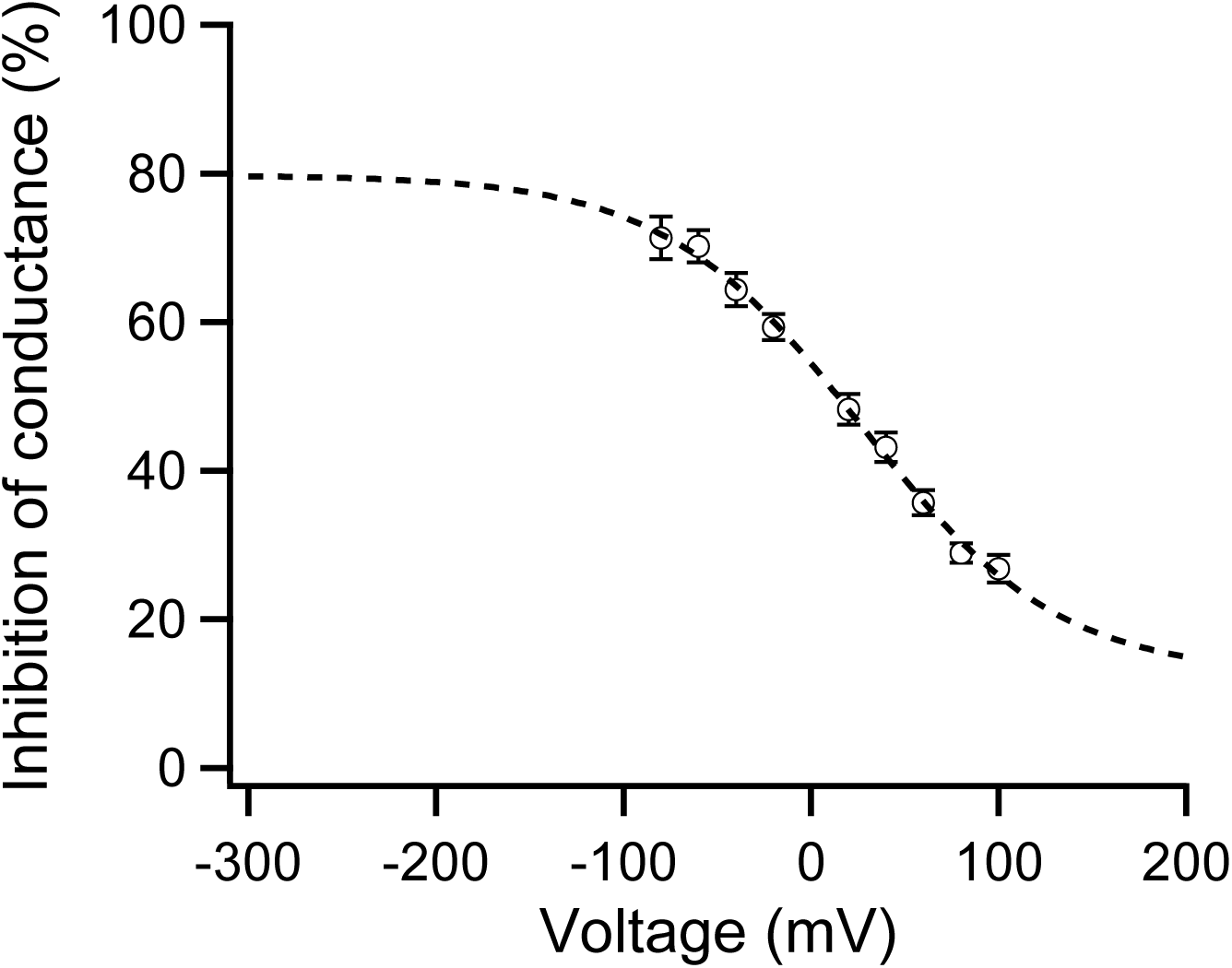
Voltage-dependence of proton block of TRPV1 current. This figure is adapted from our previous study^22^. Proton block is calculated from the OFF response recorded in outside-out patches. A Boltzmann function was superimposed (V_1/2_ = 26.6 mV, q_app_ = 0.48 e_0_, n = 5- to-9). All statistics are given as mean ± s.e.m.

**Extended Data Figure 5.**
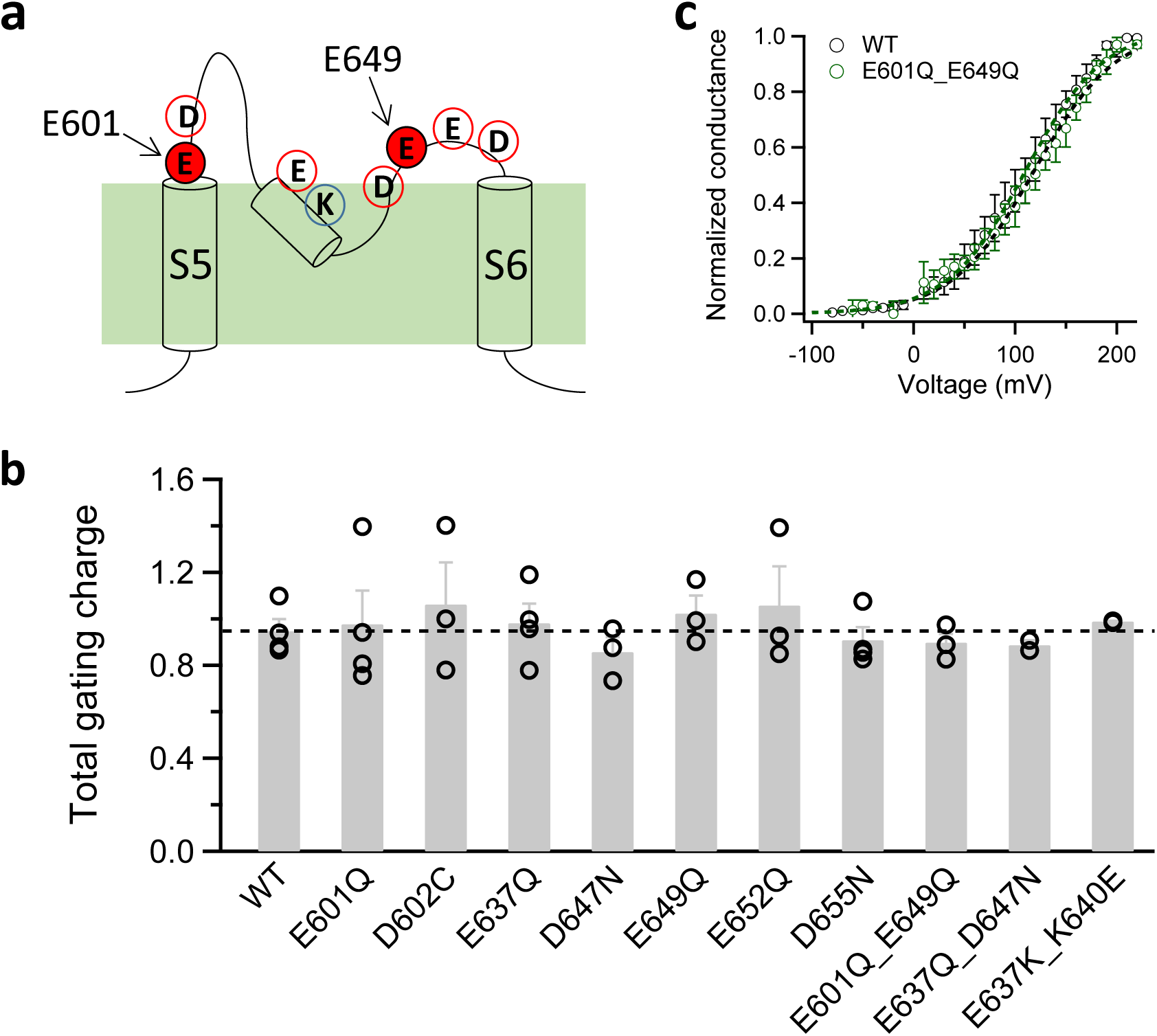
Neutralizing individual charged residues in the outer pore near transmembrane voltage field did not eliminate voltage activation in TRPV1. (**a**) A topology plot of the pore region of TRPV1 showing the charged residues near transmembrane voltage field. Two residues, E601 and E649, known to be critical for proton activation are highlighted in red. (**b**) Total gating charge measured from each single and double mutant. A dashed line in black indicate the total gating charge of WT. All data are given as mean ± s.e.m. (n = 3-to-4). (**c**) The E601Q_E649Q double mutant (green) exhibits a similar G-V curve (V_1/2_ = 114.1 ± 10.9 mV, q_app_ = 0.90 ± 0.04 e_0_, n = 3) as WT (black).

**Extended Data Figure 6.**
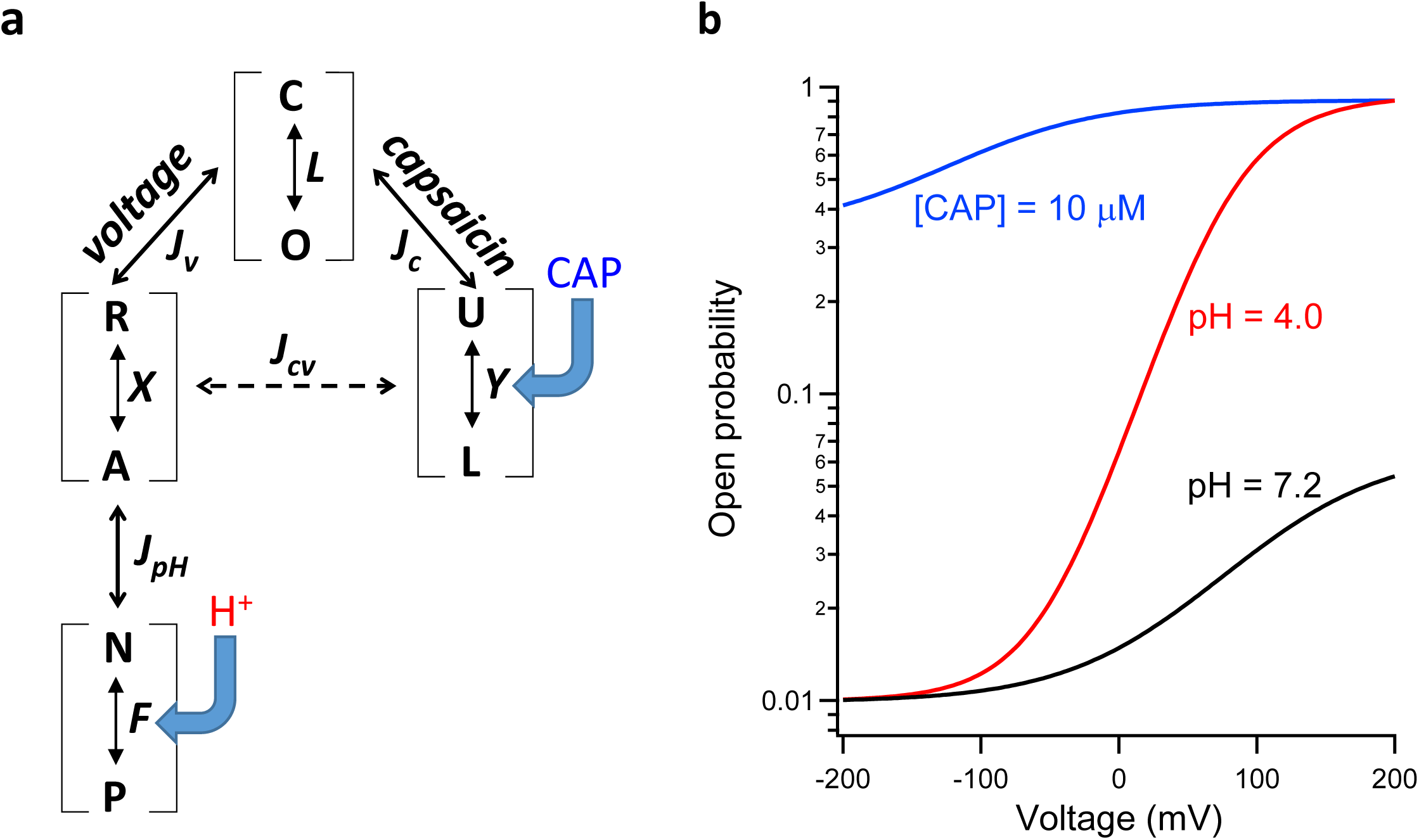
A model of multi-allosteric coupling satisfactorily predicts the behavior of TRPV1 gating by voltage (black), capsaicin (blue) and proton (red). In this model (**a**), the voltage-dependent transition R↔A is assumed to be completely separated from the proton-driven transition N↔P, as shown in Scheme III (Figure 3i), with the total gating charge movement split evenly between these transitions. Detailed description of the multi-allosteric coupling and determination of the open probability were given in our publication Cao et al. (JGP, 2014). Values for the parameters in the model that yielded the voltage-dependent Po shown in (**b**) were mostly identical to those used in Cao et al. (JGP, 2014): *L* = 0.01, *q* = 0.95 e_0_, *T* = 293.15 K, *J_V_* = 7, *V_1/2_* = 130.4 mV, *J_C_* = 150, *J_pH_* = 250, *J_CV_* = 25, [capsaicin] = 10 μM, *E* = 5 X 10^4^, *pH* = 4, *F* = 4 X 10^3^.

**Extended Data Figure 7.**
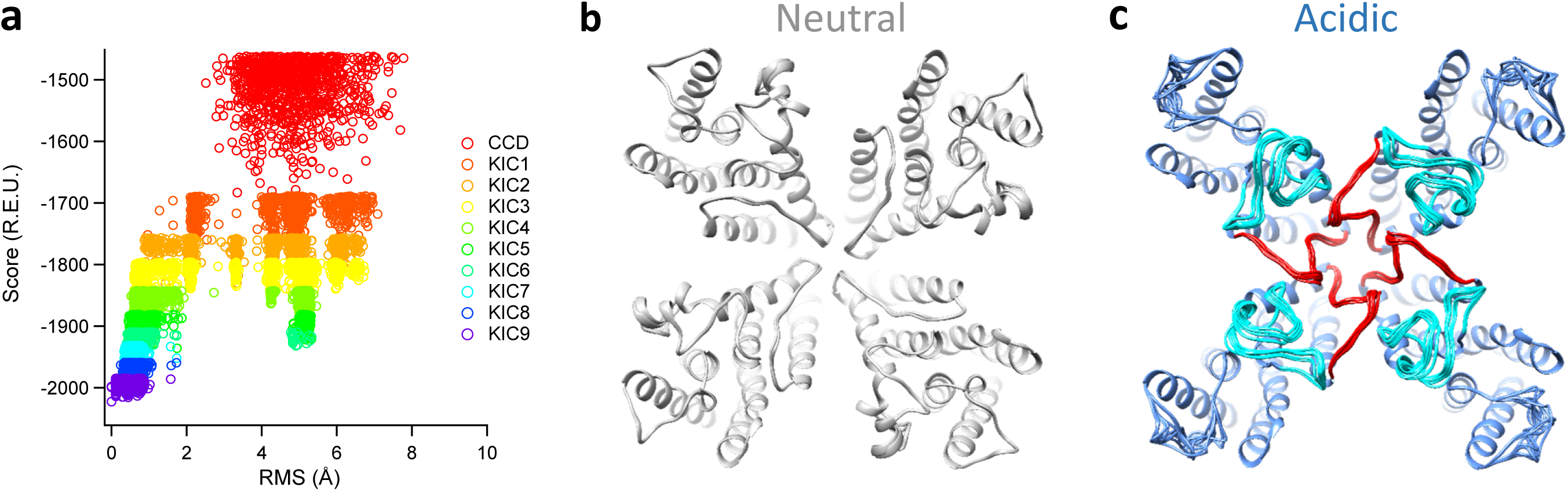
Modeling of the outer pore region of TRPV1 under neutral and acidic pH conditions. (a) Plot of the Rosetta full-atom score versus *de novo* loop RMS for ten rounds of iterative loop modeling under neutral condition. Low energy models have a more negative value of the full-atom score. (b) Overlap of the top 10 models with the lowest energy, which are well converged after iterative loop modeling under neutral condition. (c) Top 10 models with the lowest energy are well converged after iterative loop modeling under acidic (pH 4.0) condition. The turret and the linker between pore helix and S6 are colored in cyan and red, respectively.

